# Oxidative phosphorylation pathway disruption is an alternative pathological mechanism leading to Diamond-Blackfan anemia

**DOI:** 10.1101/2022.03.14.484221

**Authors:** Rudan Xiao, Lijuan Zhang, Zijuan Xin, Junwei Zhu, Qian Zhang, Siyun Chu, Jing Wu, Lu Zhang, Yang Wan, Xiaojuan Chen, Weiping Yuan, Zhaojun Zhang, Xiaofan Zhu, Xiangdong Fang

**Author notes:** Corresponding authors. **Xiangdong Fang**, Beijing Institute of Genomics & China National Center for Bioinformation, Beijing 100101, P. R. China, **Xiaofan Zhu**, State Key Laboratory of Experimental Hematology, Institute of Hematology and Blood Diseases Hospital, Chinese Academy of Medical Sciences & Peking Union Medical College, Tianjin 300020, China, **Zhaojun Zhang**, Beijing Institute of Genomics & China National Center for Bioinformation, Beijing 100101, P. R. China. R.X., L.Z., and Z.X. contributed equally to this study.

## Abstract

Diamond-Blackfan anemia (DBA) is a rare congenital disorder characterized by the failure of erythroid progenitor differentiation; however, the molecular mechanisms leading to erythroid defects remain unclear. By analyzing the transcriptomic profiles of bone marrow from patients with DBA (n = 10), we identified the dysfunction of the oxidative phosphorylation (OXPHOS) pathway as a possible cause of DBA. We established a DBA cell model using differentiating hematopoietic stem progenitor cells in which the OXPHOS pathway was suppressed to completely recapitulate the defects in erythroid progenitor differentiation, ribosome biogenesis, and heme biosynthesis, which are representative characteristics of patients with DBA. Disruption of the OXPHOS pathway led to ribosomal defects and associated erythroid defects via abolishment of the Ran GTPase activating protein RanGAP1, which is pivotal in the RNA transport pathway. The composition of the ribosomal proteins in the established DBA cells was unchanged, but an overall reduction in ribosomal protein levels was observed, leading to an alteration in the translation of a subset of transcripts specific to erythropoiesis. We revealed that the OXPHOS pathway participates in erythropoiesis, particularly at an early stage, and reinforced the relationship between the OXPHOS pathway and erythropoiesis. Coenzyme Q10, an activator of OXPHOS, largely rescued the erythroid defects in DBA cells. Our results reveal that OXPHOS repression is an alternative pathological mechanism leading to DBA, demonstrating its potential as a therapeutic pathway.

**Key Points:**

- Oxidative phosphorylation (OXPHOS) pathway disruption is an alternative pathological mechanism underlying Diamond-Blackfan anemia (DBA).
- Suppression of OXPHOS leads to defects in erythropoiesis and ribosomal biogenesis via the RanGAP1 protein.

## Introduction

Diamond-Blackfan anemia (DBA) is a rare inherited bone marrow failure syndrome characterized by erythropoiesis failure, leading to hypoplastic anemia, congenital defects, and a predisposition for malignancies [1]. Patients with DBA occasionally present with neutropenia or thrombocytosis [2] and typically present with erythroblastopenia caused by differentiation failure of only the erythroid linage. Understanding the process of erythropoiesis and its underlying mechanisms, which are currently unclear, may provide insights into the pathological mechanisms of DBA.

Gene mutations, including those in ribosomal and non-ribosomal protein genes, have been identified as the causative agents of DBA [3, 4]. Most of these mutations lead to impaired ribosome biogenesis and erythroid defects in erythroid progenitor cells because these cells are sensitive to apoptosis caused by p53 pathway activation resulting from haploinsufficiency of ribosomal proteins (RPs) in DBA [5]. Erythroid progenitor cells are also extremely sensitive to impaired translation, which may be attributable to the rapid cell division or high translational demand for globin synthesis in the erythroid cell lineage [5-7]. L-leucine potently stimulates protein synthesis by activating mTOR signaling, which underlies enhanced erythroid progenitor differentiation observed in DBA models [8, 9]. Heme biosynthesis facilitates globin production during erythropoiesis. In DBA, production of heme is higher than that of globin, and decreasing heme synthesis improves red blood cell production [10]. Impaired ribosome biosynthesis is also a major feature of DBA pathology [11, 12], which can result in loss of translation of select transcripts specifically sensitive to erythropoiesis. Previously discovered pathological mechanisms underlying DBA are closely related to various aspects of erythropoiesis. We hypothesized that identifying the events involved in the regulation of early erythropoiesis could improve our understanding of DBA pathology and may be useful in clinical applications.

Oxidative phosphorylation (OXPHOS) generates most of the energy required for cells through the mitochondrial electron transport chain [13-15]. OXPHOS is required for cell lineage specification and development [16] and is responsible for providing energy for differentiating hematopoietic stem progenitor cells (HSPCs) towards the erythroid lineage [17] but not towards other lineages [18]. OXPHOS deficiency is among the most common genetically inherited condi and can result in various diseases with an estimated prevalence of 1:5000 in humans [13]. However, the association between the OXPHOS pathway and erythropoiesis or related diseases is largely unknown.

Our study aimed to examine the role of the OXPHOS pathway in erythropoiesis and determine the pathological mechanism linking this pathway with the pathology of DBA. Our findings may improve the understanding of erythropoiesis and provide more insights into the pathology of DBA.

## Materials and Methods

### Clinical samples

Bone marrow (BM) samples from patients with DBA (n = 10) and patients with idiopathic thrombocytopenic purpura (ITP; n = 3), who were included as the healthy control group [19], were obtained from Tianjin Blood Disease Hospital, Chinese Academy of Medical Sciences & Peking Union Medical College. DBA was clinically diagnosed according to established criteria [20, 21]. The patients with DBA were aged 4–120 months (mean 25.2 months). All patients with DBA had been diagnosed based on previously reported procedures [22]. Three patients with physical deformities had mutations in RPs, including DBA-5 (*RPL11* mutation, limb deformities), DBA-9 (*RPS19* mutation), and DBA-10 (*RPL5* mutation, finger deformities) (Supplemental Table 1). Human cord blood samples from full-term newborns were obtained from Beijing Obstetrics and Gynecology Hospital, Capital Medical University (Beijing, China). The subjects provided written informed consent and the study was performed in accordance with the Declaration of Helsinki. The protocols for sample collection and use were approved by the ethics committee of the Beijing Institute of Genomics, Chinese Academy of Sciences (approval no. 2017H006).

### CD34+ cells

We isolated CD34+ cells from human cord blood and BM samples using an EasySep™ Human CD34 Positive Selection Kit II (STEMCELL Technologies, Vancouver, Canada); sample purity of >95% was confirmed using flow cytometry. When an expansion phase was required, CD34+ cells were cultured in StemSpan SFEM II medium (STEMCELL Technologies) supplemented with 50 μg/mL stem cell factor, 50 μg/mL Flt3 ligand (PeproTech, Rocky Hill, NJ, USA), 50 μg/mL thrombopoietin (PeproTech), and 1% penicillin and streptomycin solution for 6 days prior to differentiation. CD34+ cell purity was >95% after expansion, as determined using fluorescence-activated cell sorting. Erythroid cells were differentiated as described previously [23] with incubation at 37°C and 5% CO_2_.

### DBA cell model

Differentiating HSPCs in which the OXPHOS pathway was suppressed towards the erythroid lineage were used as the primary cell model to explore the relationship between OXPHOS and DBA pathology. HSPCs were isolated from human cord blood or BM samples from healthy control patients. Rotenone (Rot, R817233, Sigma-Aldrich, St. Louis, MO, USA), which inhibits electron transfer from the iron-sulfur center to ubiquinone, thereby inhibiting ATP production, was added at 0, 0.5, and 1 µM concentrations to the culture medium when erythroid differentiation was initiated from CD34+ cells. OXPHOS inhibition was measured using flow cytometry at 24, 48, and 72 h, at the optimized concentration of Rot (1 µM). Coenzyme Q10 (CoQ10, C111044, Sigma-Aldrich), which activates OXPHOS by transporting electrons to the electron transport chain during respiration, was used to rescue the inhibitory effect of Rot on OXPHOS. CoQ10 (25 and 50 μM) was added to the culture medium for an additional 72 h after Rot treatment, and the difference in cell membrane potential was measured using flow cytometry. Further, colony-forming unit assay was performed as previously described [23].

### Flow cytometry

To detect OXPHOS inhibition, Rot-treated cells were collected and centrifuged at 300 ×*g* for 10 min, and the pellet was resuspended in 70 µL Hanks’ buffer; 20 µL of this cell suspension was retained without staining as the blank control. When a change in mitochondrial membrane potential was detected, 10 μM rhodamine 123 fluorescent dye (R8004, Sigma-Aldrich) was added to the 50 µL cell suspension, which was then incubated at 37°C for 30 min in the dark. The cells were washed twice with Hanks’ buffer, and flow cytometry was performed.

To detect cell-surface markers of the differentiated cells, flow cytometry was performed as per a previous study [23]. Fluorescence-activated cell sorting analysis was conducted using a BD Biosciences Canto II flow cytometer (BD Biosciences, Franklin Lakes, NJ, USA). Data were analyzed using FlowJo 10.0.7 software (TreeStar, Ashland, OR, USA).

### Immunofluorescence assay

On day 7, 10^5^–10^7^ differentiated cells were collected into 2 mL centrifuge tubes, gently mixed with 2 mL of Dulbecco’s phosphate-buffered saline (DPBS), and centrifuged at 1000 ×*g* for 5 min. After washing the cell pellet twice with DPBS, 4% paraformaldehyde was immediately added to fix the cells for 10 min, followed by centrifugation at 1000 ×*g* for 5 min. The pellet was washed thrice with DPBS and then 0.4% Triton-X-100 was added for 15 min, before centrifuging the cells at 1000 ×*g* for 5 min. The pellet was washed with DPBS thrice, and 5% bovine serum albumin (BSA)-DPBS was added. After centrifugation at 1000 ×*g* for 5 min, primary antibodies against RPS6 (ab80158, Abcam, Cambridge, UK) and RPS3 (ab128995, Abcam) were separately added to the pelleted cells, which were allowed to stand at 25°C for 2 h. After washing with DPBS thrice, fluorescently labeled goat anti-rabbit secondary antibody was added, incubated at room temperature for 1 h, and washed thrice with DPBS. The cells were evenly spread and dried on clean glass slides. DAPI was added, anti-fluorescence quencher coverslips were mounted, and images were photographed using an upright fluorescence microscope (LSM710, Zeiss, Germany).

### Knockdown of Ran GTPase activating protein (RanGAP1)

Plasmid preparation and procedures for knocking down RanGAP1 were as previously reported [23]. Primer sequences are shown in Supplemental Table 2.

### Real-time PCR

Total RNA extraction and real-time PCR were carried out as previously described [24] using the primer sequences shown in Supplemental Table 2.

### Western blotting

Cells were lysed in RIPA buffer (50 mM Tris-HCl at pH 7.4, 150 mM NaCl, 0.1% SDS, 1% NP-40, 0.25% sodium deoxycholate, 1 mM dithiothreitol) supplemented with protease inhibitors (Roche, Basel, Switzerland) and incubated for 30 min on ice. After centrifugation at 18,800 ×*g* for 10 min, the protein concentration in the supernatant was determined. Equal amounts of protein were separated using sodium dodecyl sulfate (SDS)-polyacrylamide gel electrophoresis in a Tris/glycine/SDS running buffer. The separated proteins were transferred onto a polyvinylidene fluoride membrane in Tris/glycine transfer buffer, which was then blocked with 3% BSA-TBST (Tris buffered saline with Tween 20) for 1 h at room temperature. The membranes were probed with primary RanGAP1 mouse monoclonal antibody (ab92360, Abcam) at a 1:500 dilution and HBG2 (ab137096, Abcam), GATA1 (ab181544, Abcam), and β-actin mouse monoclonal antibodies (HC201-01, TransGen Biotech, Beijing, China) at a 1:1000 dilution in 3% BSA-TBST at 4°C overnight. The membranes were subsequently rinsed with TBST thrice for 10 min each and incubated with secondary antibody for 1 h at room temperature. Finally, the membranes were rinsed with TBST solution thrice and imaged using an Image Quant ECL machine (Tanon 5200, Tanon, Shanghai, China).

### RNA-sequencing

BM samples were lysed to remove red blood cells, and mononuclear cells were isolated using a Ficoll density gradient, followed by extraction of total RNA using TRIzol for RNA-sequencing (RNA-Seq) library construction. Total RNA was isolated from differentiated cells on days 3 and 7 (two biological replicates each). RNA-Seq libraries were sequenced on an Illumina sequencing platform (San Diego, CA, USA), generating 150 bp paired-end reads. At least 4G of data were collected for further analysis. Quality control was analyzed using FastQC software, and the original sequence FASTQ file was compared to the UCSC version hg38 reference file using HISAT2 software. The gene expression matrix file obtained using StringTie software was used as the input file to analyze differentially expressed genes (DEGs). The R package edgeR was used for DEG analysis with a biological coefficient of variation of 0.1. The DEG screening criteria were |log fold-change (FC)| > 1 and *p* < 0.05. DEGs were used for bioinformatics analysis, including Kyoto Encyclopedia of Genes and Genomes (KEGG), Gene Ontology (GO), and Gene Set Enrichment Analysis (GSEA).

### Single-cell transcriptome sequencing data analysis

Single-cell RNA-seq data were downloaded from the Gene Expression Omnibus database (GSE139369 and GSE150774). Transcriptome data of 49,707 single cells were collected, including 12,602 BM mononuclear cells, 8,176 CD34+-enriched BM mononuclear cells, 14,804 peripheral blood mononuclear cells, and 14,125 terminal erythroid cells (CD34-CD235a+) isolated directly from adult BM. These data were generated using 10× Genomics 3 single-cell RNA sequencing. Gene expression was standardized using the default parameters of R package Seurat (version 4.0.6). Principal component analysis (dims = 20) was performed to reduce the dimensions of the data. UMAP was used to visualize individual cells. We annotated the cells using the R package SingleR (version 1.8.0) and published datasets [25, 26]. Cells at different stages of hematopoietic stem cell (HSC) differentiation into red blood cells were selected, and hallmark gene sets were downloaded from the GSEA database. GSEA was analyzed using the R package GSVA (version 1.14.1) to determine the OXPHOS gene set enrichment scores in erythroid cells at different stages of differentiation.

### Proteome analysis

Differentiated cells on day 7 were used for mass spectrometry (MS) and RNA-Seq analyses (three biological replicates each). Protein extraction, trypsin digestion, tandem mass tag (TMT) labeling of peptides, and liquid chromatography with tandem MS (LC-MS/MS) analysis were carried out as previously described [27]. The resulting spectra were individually analyzed against the database using Proteome Discoverer 2.2 (PD 2.2, Thermo Fisher Scientific). The mass tolerance of the precursor ion was 10 ppm and that of the production was 0.02 Da. No more than two miscleavage sites were allowed. The identified protein contained at least one unique peptide with a false discovery rate (FDR) ≤ 1.0%. Reporter quantification (TMT 10-plex) was used for TMT quantification. Significantly differentially expressed proteins between the experimental and control groups were screened using the following criterion: *p* < 0.05 and |log2FC| > 0. GO and KEGG analyses were performed using ClusterProfiler (version 4.0.2).

### Data availability

The supporting RNA-Seq datasets for the DBA samples and differentiated cells are respectively available from the Genome Sequence Archive with accession numbers HRA002012 and HRA002000. The following link was created to allow a review of the records: https://ngdc.cncb.ac.cn/gsa-human/. The supporting proteomics datasets are available from the Genome Sequence Archive with accession number OMIX971; the following link was created to allow a review of the record: https://ngdc.cncb.ac.cn/omix/.

### Statistical analysis

Data are presented as mean ± standard deviation (SD). Statistical analyses were performed using GraphPad Prism (version 8.0.2) and *p* < 0.05 was considered statistically significant. Student’s t-test was used and significance indicated as follows: \**p* < 0.05; \*\**p* < 0.01; \*\*\**p* < 0.001.

## Results

### OXPHOS disruption may cause DBA

Clinical characteristics of the DBA cohort are shown in Supplemental Table 1. To explore the pathogenesis of DBA from a broad perspective, we subjected the BM samples of the DBA and control groups to bulk transcriptome sequencing. Analysis revealed that abnormally disturbed genes in the DBA samples were significantly enriched in the OXPHOS pathway (*p* = 2.51E-41), mitochondrial dysfunction (*p* = 1.99E-41), mTOR pathway (*p* = 6.76E-5), and heme biosynthesis II (*p* = 1.10E-4) (Figure 1A). The OXPHOS pathway is highly dependent on the proteins encoded by the mitochondrial genome, which were not reported in DBA. Analysis of the transcriptome profiles of DBA samples against the control showed that OXPHOS was inhibited in 7 of 10 DBA cases, including the case with an *RPL11* mutation. Furthermore, clustering of the

**Figure 1.**
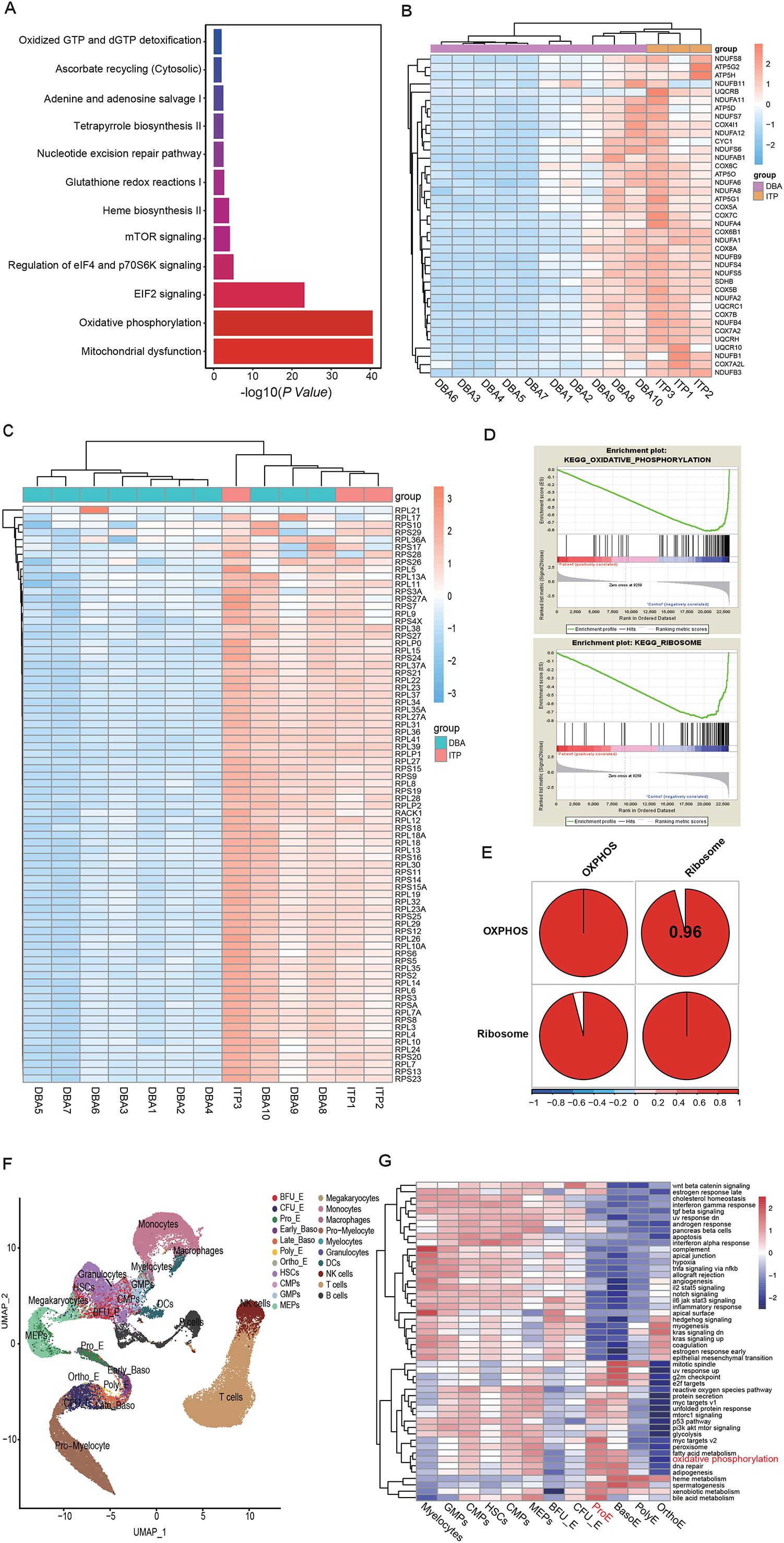
Association of disrupted OXPHOS pathway with DBA pathology. A. Pathway enrichment analysis of DEGs in DBA samples against control samples showing that the OXPHOS pathway was significantly suppressed in patients with DBA. B. Heatmap analysis of key genes in the OXPHOS pathway mapped to all DBA and control samples showing that the OXPHOS pathway was suppressed in 7 of 10 patients with DBA. Among the DBA patients, patient 5 (*RPL11* mutation) showed suppression of the OXPHOS pathway, whereas DBA patients 9 (*RPS19* mutation) and 10 (*RPL5* mutation) did not. C. Heatmap analysis of genes in the ribosome biogenesis pathway against all DBA and control samples showing that the ribosome biogenesis pathway was also suppressed in the same batch of DBA samples (7 of 10) as in those with the suppressed OXPHOS pathway. D. GSEA analysis showing enrichment of suppressed OXPHOS and the ribosome pathway in DBA samples compared to in controls. E. Correlation analysis of OXPHOS and the ribosome biogenesis pathway in patients with DBA. After calculating the TPM expression value of OXPHOS and ribosome pathway genes in each DBA sample, we standardized the gene expression value relative to the min-max, mapped the data to the [0,1] interval, and used the R package “ssGSEA” to analyze the OXPHOS and ribosome pathways and determine the score matrix of these two pathways in different samples. R package “corrplot” was used to calculate the correlation of gene expression of these two pathways based on the score matrix. F. UMAP plot of single-cell cluster distribution of cells in the bone marrow from healthy individuals. This plot covers all differentiated cells towards the erythroid lineage; G. GSVA of cells at different stages of HSC differentiation into red blood cells.

DBA cases closely resembled the pattern observed for all genes in the OXPHOS pathway (Figure 1B), suggesting that OXPHOS is inhibited in DBA. Expression of genes associated with the ribosome biogenesis pathway was also significantly inhibited in DBA cases compared with the control (Figure 1C), and GSEA analysis showed that genes associated with these two suppressed pathways were enriched in DBA samples (Figure 1D). The transcriptome profiles of OXPHOS and the ribosome pathway in DBA were highly positively correlated (up to 0.96) (Figure 1E), indicating that OXPHOS and ribosome biogenesis may utilize a mutual regulatory mechanism in DBA. As DBA is a ribosomopathy [28, 29], suppression of the OXPHOS pathway may be an alternative pathogenic pathway of DBA.

Further, to explore the association of the OXPHOS pathway with erythropoiesis, we analyzed single-cell transcriptome data using the BM of healthy individuals and profiled cells differentiated towards the erythroid lineage under normal physiological conditions (Figure 1F). The OXPHOS pathway was finely controlled throughout erythropoiesis, exhibiting the highest activity at the pro-erythroblast stage. This stage is thought to demand higher energy for the stage transition from burst-forming unit-erythroid (BFU-E) / colony-forming unit-erythroid (CFU-E) to the early stage of terminal erythroid differentiation, suggesting that suppression of OXPHOS blocks erythroid differentiation at the progenitor cell stage (Figure 1G). We also identified other interesting pathways regulating erythropoiesis, such as heme biosynthesis, which is specifically active throughout terminal erythroid differentiation and is initiated at the pro-erythroblast stage; these results are consistent with the pathology of DBA [10].

### OXPHOS suppression leads to failure of erythroid progenitor differentiation

The development of HSPCs into erythroid cells can serve as a cell model for exploring the pathogenesis of hematological disorders associated with mitochondrial dysregulation [18]. We developed an *ex vivo* culture system to differentiate human HSPCs into erythroid cells in which the OXPHOS pathway is inhibited by Rot [30]. A greater reduction in mitochondrial membrane potential (P3) indicates greater suppression of the OXPHOS pathway [31]. Rot inhibited OXPHOS in a dose- and time-dependent manner (Figure 2A, B) and decreased ATP production (data not shown); thus, we developed a cell model of DBA cells in which the OXPHOS pathway was suppressed.

**Figure 2.**
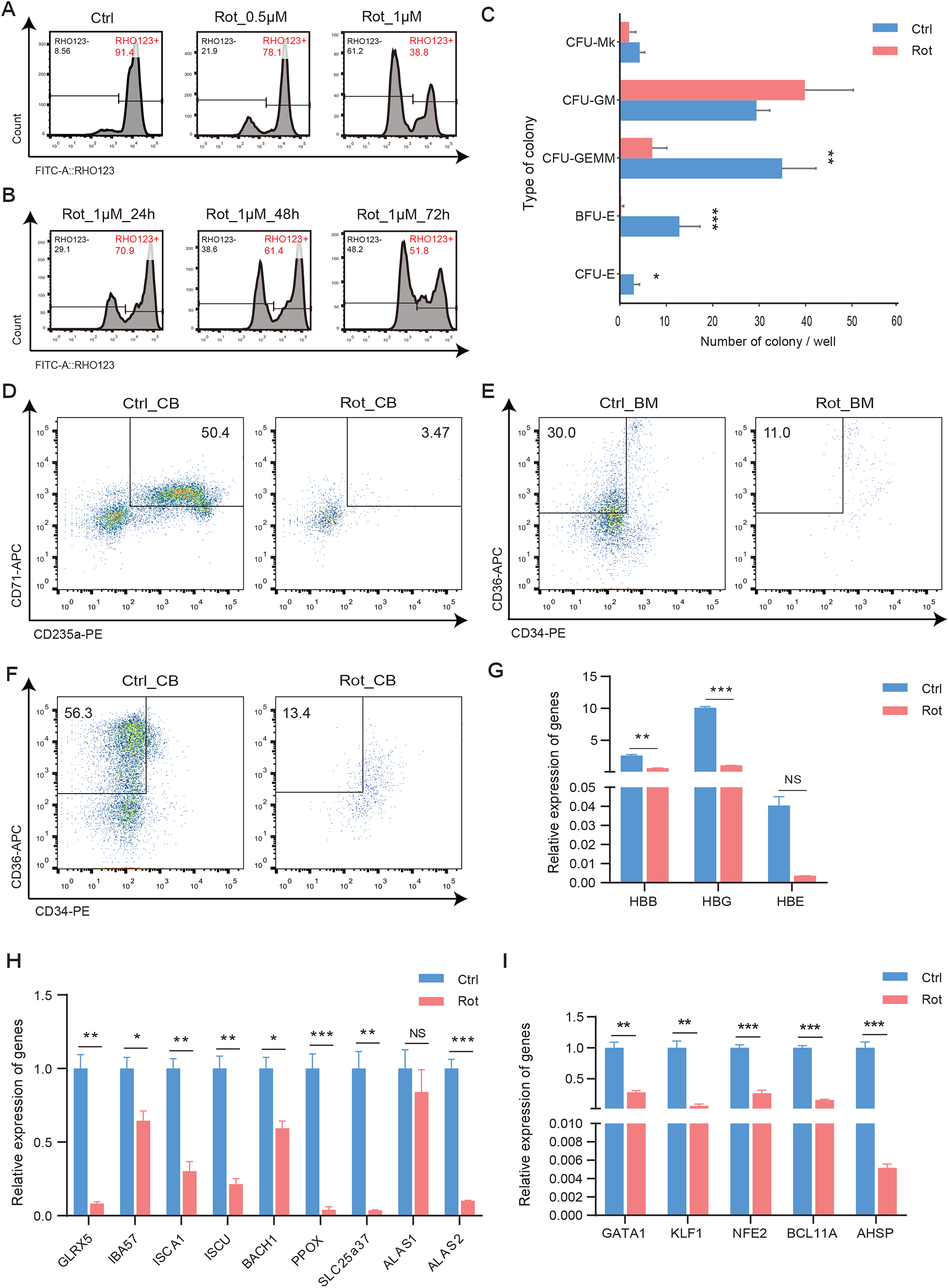
Suppression of OXPHOS specifically leads to failure of erythroid progenitor differentiation. A–B. Flow cytometry analysis of Rot-treated HSPCs showing inhibition of the OXPHOS pathway by Rot in a dose- and time-dependent manner, respectively. The level of OXPHOS inhibition was proportional to the ratio of the left peak count to the total count. C. Colony-forming potential analysis of Rot-treated HSPCs showing specific erythroid defects that were not observed in other lineages. Triple replicates were set for each treatment with or without Rot in the 96-well plate and data represent the mean ± SD. D. Flow cytometry analysis showing a significantly reduced number of CD71+GlyA+ double-positive erythroid cells differentiated from Rot-treated HSPCs compared with control cells. E–F. Flow cytometry analysis showing severe blockage at the stage of erythroid progenitors (CD34+CD36+) derived from Rot-treated HSPCs in healthy BM and cord blood in differentiating DBA cells on day 7. G–I. Real-time PCR analysis showing a dramatic decrease in the expression of erythroid-specific globin genes (*HBG, HBB*), heme biogenesis-related genes, and other erythroid-specific genes including transcription factors in DBA cells on day 7. *ACTIN* was used as reference gene in real-time PCR assays. Student’s *t*-test was used in this study: \**p* < 0.05; \*\**p* < 0.01; \*\*\**p* < 0.001.

Blocked differentiation at the BFU-E or CFU-E stages is a hallmark of DBA [10, 32]; if suppression of OXPHOS causes DBA, erythropoiesis failure is also expected to occur. To evaluate this hypothesis, we conducted a CFU assay with CD34+ cells as the starting cells in which OXPHOS was inhibited. DBA cells with OXPHOS inhibition generated normal numbers of myeloid cells but significantly reduced colony numbers of BFU-E, CFU-E, and CFU-GEMM, which include erythroid lineage cells, indicating specific loss of erythroid progenitors (Figure 2C). This indicates that erythropoiesis was impaired at the same stage as in patients with DBA. Flow cytometry results showed that inhibition of OXPHOS led to a visible decrease in differentiated CD71+GlyA+ erythroid cells (3.57%), which was caused by obvious blockage at the stage of erythroid progenitors derived from both BM and cord blood sources (Figure 2D–F). Additionally, erythroid progenitors (CD71+ GlyA-) showed a decreasing trend beginning on day 3 in both BM and cord blood sources (Supplemental Figure 1 A-B). Decreased globin expression is characteristic of DBA phenotypes [10]. Globin expression showed a differential pattern in differentiated erythroid cells with or without Rot treatment on day 7, with dramatically reduced expression of *HBB* and *HBG. HBG* showed the highest expression among the globins (Figure 2G), indicating that erythroid defects in DBA cells occur at the early stage of erythroblasts or before entering the pro-erythroblast stage. Reduced globin expression in DBA cells may result from disruption of heme production [10]. The expression of most genes involved in heme biosynthesis, particularly *ALAS2*, which is the initial and rate-limiting gene of heme biosynthesis, gradually decreased in DBA cells during erythroid differentiation (Figure 2H, Supplementary Figure 1C), suggesting that the disruption of heme biosynthesis is caused by suppression of OXPHOS in DBA cells. Expression of erythroid-specific genes, mostly transcription factors, were decreased in DBA cells (Figure 2I, Supplementary Figure 1D), among which *GATA1* is a DBA-causing non-RP gene [7, 33, 34]. Inhibition of OXPHOS suppression of these erythroid-specific factors reflects the failure of erythropoiesis in DBA cells. Moreover, we observed increased apoptosis and expression of apoptosis marker genes during erythroid differentiation (Supplementary Figure 1E–F), which are typical features of DBA cells [35].

### OXPHOS suppression leads to defects in the ribosome biogenesis pathway

Ribosomal abnormalities can result from defects in DBA-causative genes [35, 36] or the reduced abundance of RPs [11]. We evaluated whether alterations in RP genes occurred in the DBA cells. We comprehensively characterized the transcriptomic features of the DBA cells on days 3 and 7, which mainly represent erythroid progenitors, and further differentiated erythroid cells during HSPCs undergoing commitment to the erythroid lineage [25, 26] (Figure 3A). Significantly suppressed DEGs in DBA cells on days 3 and 7 were enriched in the OXPHOS pathway and ribosome biogenesis pathway, which are disturbed in patients with DBA (Figure 3B), suggesting that disruption of the ribosome pathway resulted from the suppression of OXPHOS. Ribosomal genes most relevant to DBA, including *RPL* and *RPS*, were significantly suppressed (*p* < 0.001) in the Rot-treated group than in the control on day 3 (Supplemental Figure 2A) and day 7 (Figure 3C). RNA transport and spliceosome pathways were also disturbed in DBA cells (Figure 3B).

**Figure 3.**
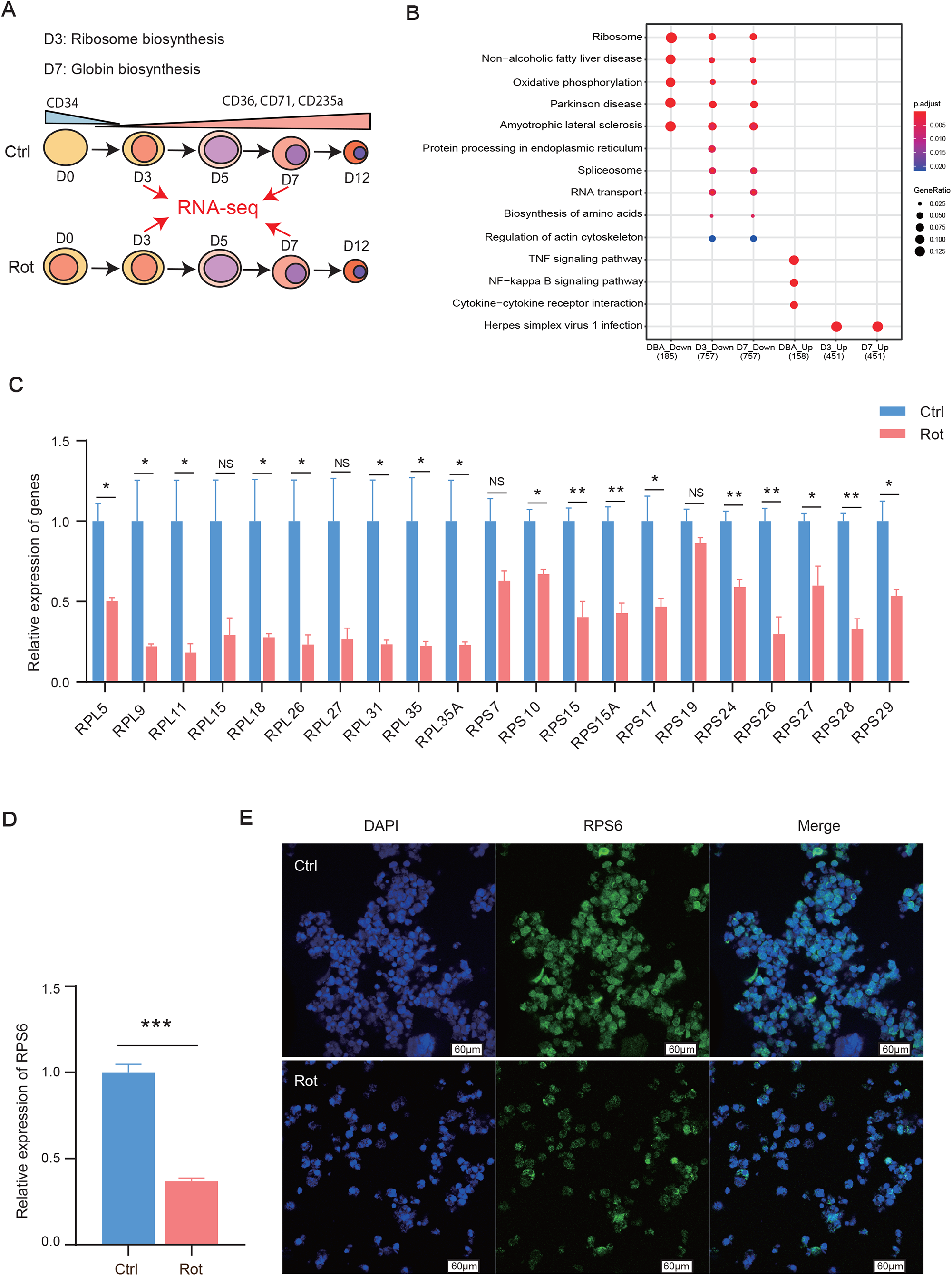
Suppression of OXPHOS leads to ribosomal defects. A. Schematic of transcriptome analysis of DBA cells during erythroid differentiation to explore the effects of inhibition of OXPHOS. Differentiating cells on days 3 and 7 treated with Rot were collected and evaluated using bulk transcriptome sequencing. The differentiated cells on days 3 and 7, respectively, represent most erythroid progenitors and earlier stage erythroblasts (e.g., Pro-E, Baso-E) during erythroid differentiation of HSPCs. B. Enrichment analysis of GO terms enriched by up- and downregulated DEGs in DBA samples as well as differentiating cells on days 3 and 7, respectively. C. Real-time PCR analysis showing significant overall reduction in the expression of most DBA causative genes identified in differentiating DBA cells on day *7. ACTIN* was used as the reference gene. D. Real-time PCR analysis showing decreased expression of *RPS6* in differentiating DBA cells on day 7. *ACTIN* was used as a reference gene. \*\*\**p* < 0.001. E. Immunofluorescence analysis of RPS6 showing defects in ribosome levels in differentiating DBA cells on day 7.

The immunofluorescence assay revealed that compared to control cells, DBA cells had significantly lower expression of RPS3 and RPS6, which are non-DBA causative proteins and specific ribosome markers, facilitating recognition of ribosome morphology and integrity [34] (Figure 3D–E, Supplemental Figure 2B– C). Thus, ribosome morphology was disrupted in DBA cells with suppressed OXPHOS, suggesting that OXPHOS is disrupted in patients with DBA, leading to defective ribosome biogenesis and decreased abundance of RPs.

### OXPHOS suppression causes ribosomal defects via RanGAP1

The RNA transport pathway was also disturbed in DBA cells (*p* < 1.00E-6) (Figure 3B), indicating that an RNA transport defect can occur with the onset of OXPHOS suppression. To explore the mechanism by which OXPHOS suppression leads to ribosome abnormalities in DBA cells, the transcriptomic data were mapped to the hsa03013-RNA transport pathway [37]. The results revealed the suppression of RAN (Ran GTPase) (Figure 4A), which is bound to GTP in the nucleus and GDP in the cytoplasm, and is essential for the translocation of RNA and proteins through the nuclear pore complex [38]. RanGAP1 associates with the nuclear pore complex, participates in regulating nuclear transport [39], and plays an important role in regulation of the RanGTP/GDP cycle [40]. We predicted that Ran-dependent nuclear transport, specifically the Ran cycle, was disrupted, which may have disturbed the expression of RanGAP1. The Rot-treated group showed abolishment of RanGAP1 (Figure 4B), whereas the SUMO1/RanGAP1 complex, which is involved in mitotic spindle formation but not nuclear transport during interphase, was only slightly suppressed [41]. Collectively, OXPHOS suppression in DBA cells may lead to defects in the ribosome via RanGAP1, thereby disrupting the Ran cycle and blocking rRNA transport.

**Figure 4.**
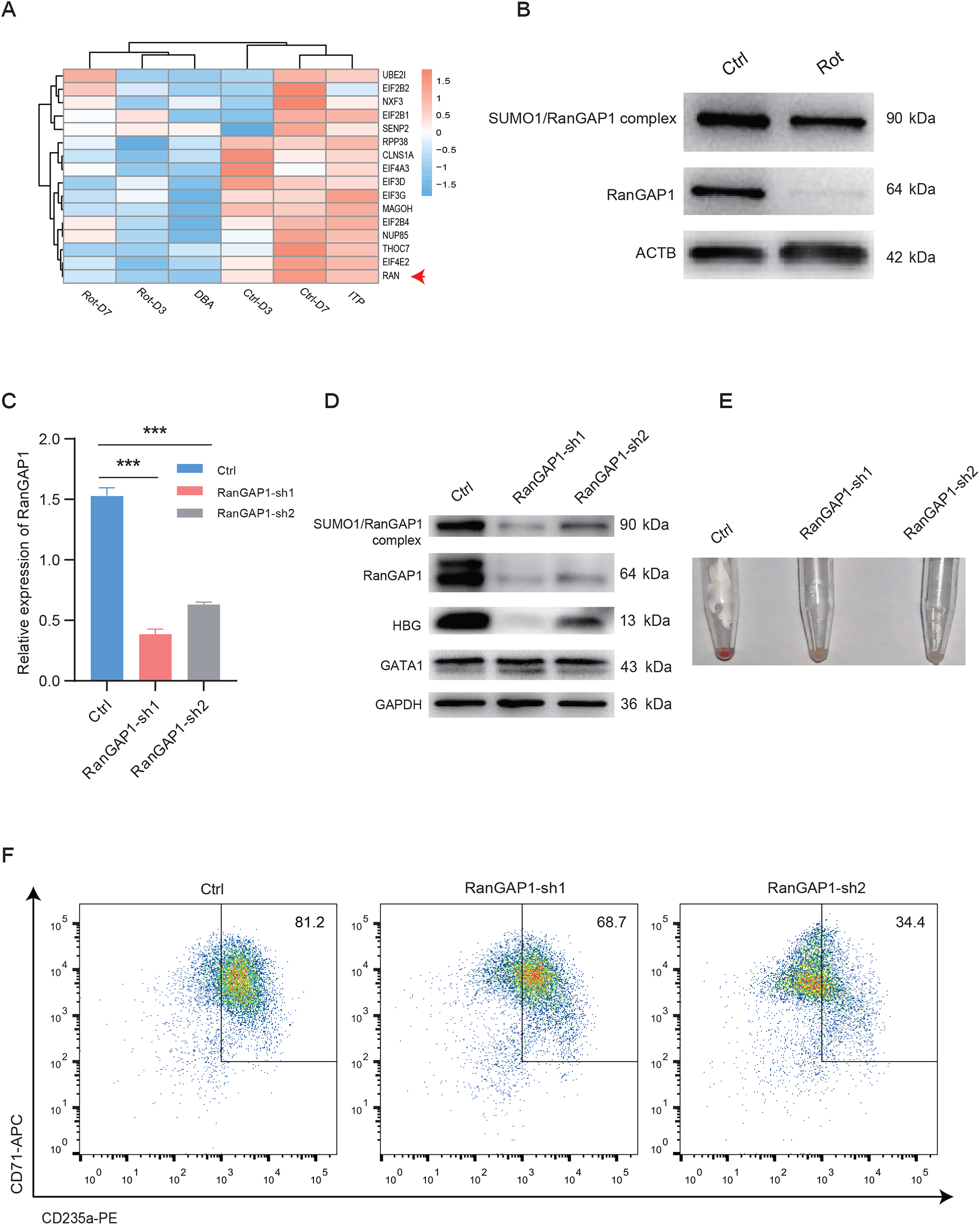
Knockdown of RanGAP1 leads to erythroid defects. A. Mapping analysis showing downregulation of RNA-transport pathway genes in patients with DBA and in DBA cells on days 3 and 7. The arrow indicates the suppressed RAN protein, which plays a key role in RNA nuclear transport and can be activated by RanGAP1. B. Western blotting assay showing the abolishment of RanGAP1 protein expression in DBA cells on day 7. Lysates for differentiated cells with or without Rot treatment were used for this experiment. The expression of SUMO1/RanGAP1 complex was simultaneously detected using the RanGAP1 antibody. Actin was used as a loading control. C. Real-time PCR analysis showing the knockdown of RanGAP1 by two independent shRNAs targeting different coding regions in RanGAP1 on day 7. \*\*\**p* < 0.001. D. Western blotting assay confirming the knockdown of RanGAP1 with shRNAs in differentiated erythroid cells from cord blood HSPCs on day 7. HBG protein expression was decreased in differentiated cells with RanGAP1 knockdown. GAPDH was used as a loading control. E. RanGAP1 knockdown delays erythroid differentiation as revealed by the color of the cell pellets on day 7. Differentiated cells containing the control vector appeared as a red pellet, whereas pelleted cells with RanGAP1 knockdown did not. E. Flow cytometry analysis showing decreased number of CD71+GlyA+ double-positive erythroid cells differentiated from cord blood HSPCs with RanGAP1 inhibition on day 7.

To test the hypothesis that RanGAP1 is a downstream target of OXPHOS suppression and leads to ribosomal defects, thereby participating in regulating erythropoiesis, we designed independent shRNAs targeting different regions of RanGAP1 (Figure 4C) and observed consistent erythroid defects in various assays. Suppression of RanGAP1 in HSPCs almost abolished HBG expression (Figure 4D) and delayed erythroid differentiation (Figure 4E), as demonstrated by the reduced number of CD71+GlyA+ double-positive cells on day 7 (Figure 1F). These defective phenotypes persisted until day 14 (Supplemental Figure 3A–E), although the expression of RanGAP1 recovered. Decreased expression of the representative DBA causative genes *RPL5* and *RPS19* may represent ribosomal defects resulting from the effects of RanGAP1 (Supplementary Figure 3F). Overall, suppression of the OXPHOS pathway may lead to ribosomal defects and associated defects in erythropoiesis via RanGAP1.

### Overall reduction in abundance of RPs and erythroid defects in DBA cells

Because ribosomal defects were observed under suppressed OXPHOS in DBA cells, we further examined the alterations in RPs and effects of reduced ribosome levels on the translation of mRNAs in DBA cells. TMT-based quantitative proteomics in parallel with transcriptome sequencing in DBA cells (Figure 5A) revealed observed perturbations of RPs at both the mRNA and protein level (Figure 5B). In the proteomic analysis, all 80 RPs were detectable by two or more unique peptides, and the estimated protein abundance was robust across biological replicates (data not shown). Ribosomal defects in DBA cells were detected as decreased expression of all RPs (Figure 5B); however, the composition of RPs was unchanged. In accordance with the finding that DBA can result from reduced ribosome abundance, but not from changes in the ribosome composition [11], we observed significant alterations in over 90% of the DBA-causative RPs that lead to erythroid defects in DBA cells.

**Figure 5.**
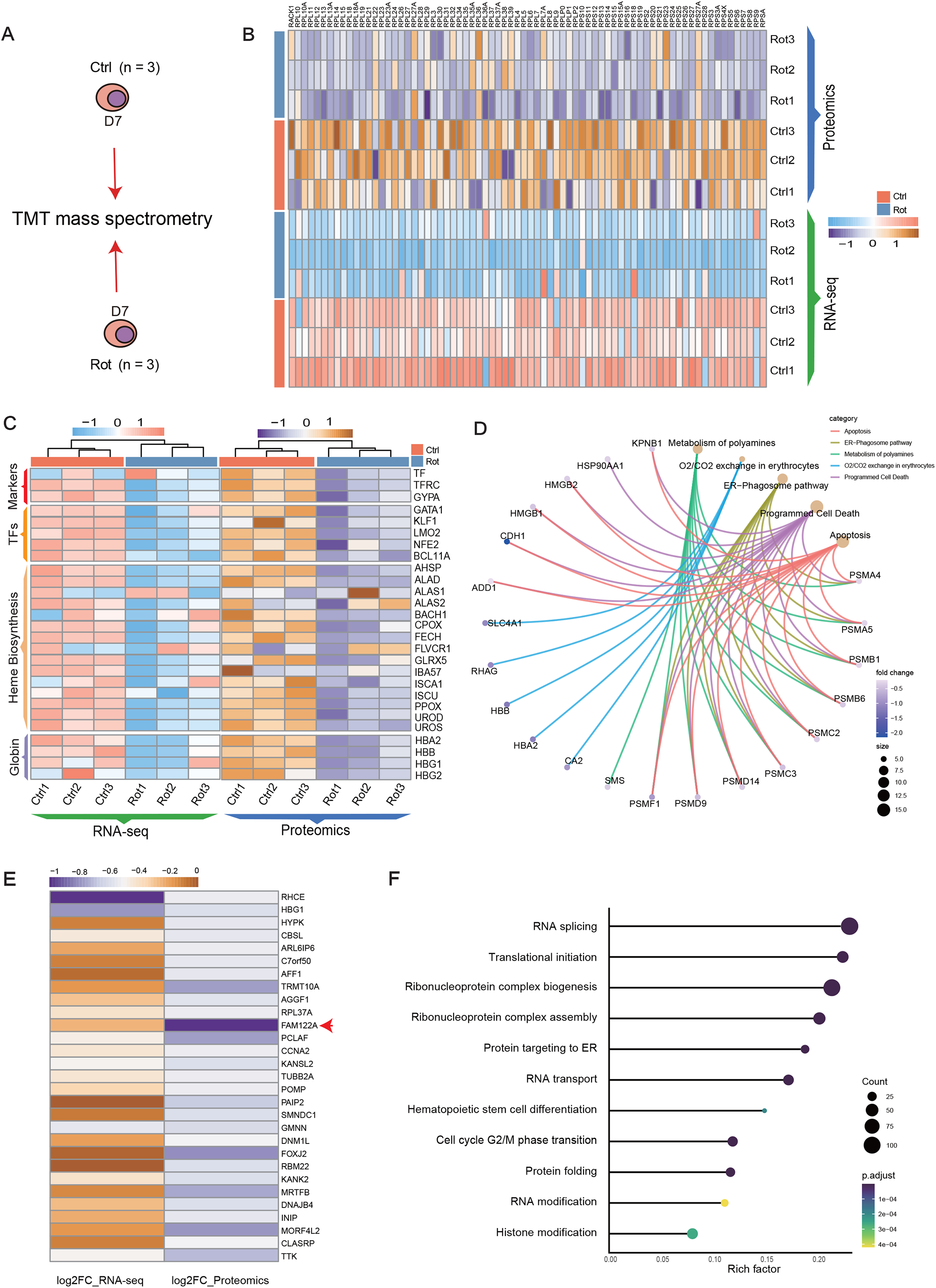
Integrative analysis showing both erythroid defects and ribosomal abnormality in DBA cells. A. Schematic of integrative analysis of proteome and transcriptome sequencing data in DBA cells. The differentiated cells on day 7, with or without Rot treatment, were simultaneously subjected to TMT mass spectrometry and bulk transcriptome sequencing. Three biological replicates were evaluated for each treatment. B. Heatmap analysis showing the overall reduction of all ribosomal constitutive proteins at both the protein and mRNA levels. C. Heatmap analysis showing erythroid defects revealed by decreased expression of erythroid-specific proteins at both the protein and mRNA levels, including erythroid markers, transcription factors, hemoglobin, and heme biosynthesis pathway-related components. D. Functional enrichment analysis of simultaneously differentially expressed genes and proteins in DBA cells. The figure was constructed using ClusterProfiler. E. Identified proteins without changes in transcript expression but with significant decreases in protein expression. The translation of these proteins may be affected by ribosome disorder resulting from inhibition of the OXPHOS pathway. The functions of most listed proteins in erythropoiesis are largely unknown. F. Functional enrichment analysis of significantly downregulated proteins without changes in transcript levels in DBA cells showing disturbed pathways. The figure was constructed using ClusterProfiler.

The high-throughput data revealed reproducible decreases in the expression of erythroid-associated markers (e.g., GYPA, CD71, TF), transcription factors (e.g., GATA1, KLF1, LMO2), and globin (e.g., HBG1, HBB, HBA2) at both the transcriptome and proteome levels (Figure 5C). Interestingly, we also observed significant suppression of the heme biosynthesis pathway in DBA cells (Figure 5C), confirming that heme biosynthesis is disrupted by inhibition of OXPHOS during the differentiation of HSPCs into erythroid cells [10, 42]. Analysis of the functions of simultaneously differentiated expressed proteins and genes revealed disturbances in O_2_/CO_2_ exchange and apoptosis in DBA cells (Figure 5D), indicating an abnormality in erythropoiesis [43]. Our integrative omics data support that defects in ribosomes and erythropoiesis occurred in DBA cells.

mRNAs are typically vulnerable to fluctuations in ribosome concentrations; thus certain cell types (e.g., erythroid cells) that depend on the protein levels of these mRNAs are sensitive to perturbations in the ribosome concentration [11, 44]. A reduction in the abundance of RPs leads to decreased translation of a subset of proteins that are sensitive to erythropoiesis [7, 35, 45]. Therefore, we comprehensively screened genes without changes in transcript expression, but showing a significant decrease in protein expression; thus, identifying inefficiently translated genes that may be sensitive to ribosomal defects in DBA cells (Figure 5E). FAM122A was the most significantly downregulated protein in DBA cells (Figure 5E) and it is also abnormally upregulated in patients with β-thalassemia [46]. Downregulation of FAM122A can lead to reduced GATA1 transcriptional activity by decreasing its association with target DNA in erythroid cells during terminal erythroid differentiation [46]. We identified a subset of sensitive genes whose functions have not been associated with ribosome defects in erythropoiesis, except for *HBG1* and *FOXJ2* [47]. These sensitive, downregulated proteins were enriched in the pathways of RNA splicing and transport, translational initiation and protein targeting to the ER, and ribonucleoprotein complex biogenesis (Figure 5F). These disturbed pathways may be associated with DBA pathology by reducing the translation of their related transcripts under the abnormal regulation of ribosomal subunits or specific RPs.

### CoQ10 rescues erythroid defects in DBA cells

To verify whether OXPHOS inhibition causes erythroid defects, DBA cells were treated with the OXPHOS activator CoQ10. The inhibitory effects of Rot on OXPHOS were significantly alleviated in CoQ10-treated DBA cells, as revealed by an increase in mitochondrial membrane potential (Figure 6A) and significant increase in ATP production compared to cells treated only with Rot (*p* < 0.001) (Figure 6B). Further, we tested the effect of CoQ10 on the expression of RanGAP1, which participates in erythropoiesis and was almost abolished by inhibition of OXPHOS in DBA cells. Interestingly, the suppression effect was completely rescued by addition of CoQ10 by reversing the inhibition of OXPHOS (Figure 6C), whereas expression of the SUMO1/GAP1 complex was not suppressed [41]. To verify whether these defects were directly caused by OXPHOS inhibition, a CoQ10 rescue experiment was performed. Flow cytometry analysis revealed that a visible decrease in differentiated GlyA+CD71+ double-positive erythroid cells (1.17%) was alleviated up to 23.4% by addition of CoQ10 (Figure 6D), suggesting that CoQ10 restores early erythroid defects in DBA cells because erythropoiesis in patients with DBA cannot progress past the proerythroblast stage [48, 49]. We also observed phenotypic changes in OXPHOS suppression in differentiated HSPCs following Giemsa staining (Figure 6E), revealing a reduced number of proerythroblasts and delayed erythroid differentiation in DBA cells, which was partially alleviated by addition of CoQ10.

**Figure 6.**
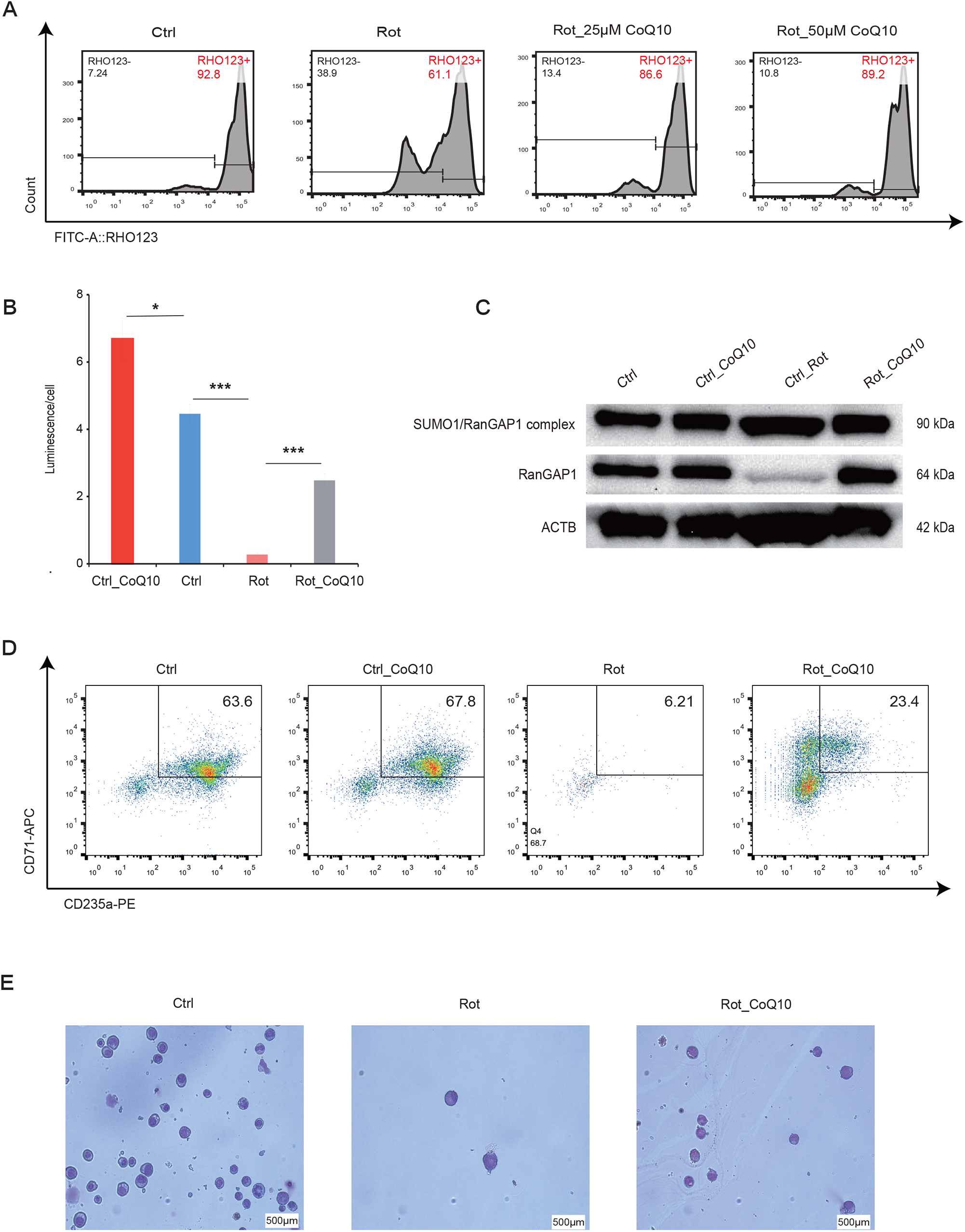
CoQ10 alleviates erythroid defects in DBA cells. A. Flow cytometry analysis showing CoQ10 largely recovers the inhibition of OXPHOS caused by Rot treatment in DBA cells. B. The ATP level was largely rescued by CoQ10 following Rot treatment in DBA cells. \**p* < 0.05; \*\*\**p* < 0.001. C. Expression of RanGAP1 in DBA cells was largely rescued by addition of CoQ10 to differentiating DBA cells on day 7. Expression of the SUMO1/RanGAP1 complex was detected using a RanGAP1 antibody. D. Flow cytometry analysis showing an increased number of CD71+GlyA+ double-positive erythroid cells following addition of CoQ10 to differentiating DBA cells on day 7. E. Giemsa staining showing that addition of CoQ10 mostly rescued erythroid defects of DBA cells. Differentiated cells were visualized on day 7.

## Discussion

The molecular mechanisms fundamental to the pathogenesis of DBA are not completely understood. We characterized the regulation of OXPHOS in erythropoiesis and explored the molecular mechanism of this pathway in the regulation of DBA pathology. We first revealed that the OXPHOS pathway is an alternative mechanism regulating DBA pathogenesis and is disturbed in many patients with DBA, with or without RP mutations. The OXPHOS pathway was not disturbed in established zebrafish DBA models with RP gene mutations [50-53], suggesting that OXPHOS is not the downstream pathway of DBA RP causative agents, but that an alternative pathogenic pathway exists in patients with DBA.

Several pathways are associated with the pathology of DBA, and RP haploinsufficiency is the primary cause of DBA. Non-RPs have also been shown to play an essential role in the pathogenesis of DBA in recent years [3, 4]. In the presence of certain RP abnormalities, pathways such as activated p53, mTOR, heme biosynthesis, and autophagy are associated with the pathology of DBA [4, 5, 10, 54-56]. Dysfunctional mitochondrial metabolism also seriously impacts erythroid differentiation. For example, loss of mitochondrial ATPase inhibitor 1 can cause severe anemia, and defects in this protein may contribute to human congenital sideroblastic anemia and other mitochondrial dysfunction-related diseases [57]. As the main energy supply system of mitochondria, OXPHOS dysfunction is crucial in the occurrence of mitochondrial diseases. The DBA cells that we established by suppressing OXPHOS recapitulated most aspects of DBA defects in erythroid progenitor differentiation, ribosome biogenesis, and heme biosynthesis. A molecular mechanism accounting for all aspects of DBA has not been determined; all mechanisms involved in DBA, including OXPHOS, are interconnected, at least partially, and none are mutually exclusive.

Disruption of OXPHOS can destroy ribosome biogenesis via the inhibition of the RNA transport pathway, leading to erythroid defects. There are two types of RNA nuclear transport, Ran-independent and Ran-dependent transport. The former is mainly mediated by the transcription–export complex, with NXF1 as the export receptor, and binding poly(A)+ RNAs [58], which are mostly mRNAs [59]. Ran-dependent export is mostly for rRNA or ribosomal subunits; thus, disruption of Ran-dependent export affects ribosome biogenesis [60, 61]. Inhibition of OXPHOS led to suppressed Ran GAP1 expression, likely affecting the Ran-dependent RNA export mechanism, resulting in rRNA transport inhibition and defects in ribosome biogenesis. We examined DBA from a new perspective and the effect of OXPHOS inhibition on the Ran cycle in HSPCs differentiating into erythroid cells. We revealed a possible connection between OXPHOS inhibition and ribosomal suppression during erythroid differentiation, providing insight into the pathophysiology of DBA.

We also examined why disruption of OXPHOS specifically leads to impaired erythropoiesis. Mitochondrial biogenesis, including oxidative phosphorylation and ATP biosynthesis, are significantly affected during the differentiation of HSPCs into erythroid-specific pro-erythroblasts [18]. In zebrafish, manipulation of the mitochondrial electron transport chain is linked to erythropoiesis [30]. The OXPHOS pathway may be conserved in some species and is directly involved in erythropoiesis. During erythroid differentiation, HSCs maintain their functions mainly via production of ATP by aerobic glycolysis, rather than by OXPHOS [17, 62-65], whereas OXPHOS provides energy for erythroid lineage specification and erythroid differentiation [17, 66]. A surge in energy is required to initiate erythroid differentiation of HSCs, during which glycolysis and OXPHOS can occur simultaneously, accompanied by a metabolic switch from glycolysis to OXPHOS, which is a driving force for erythroid differentiation [31]. OXPHOS provides energy for differentiating HSPCs towards the erythroid lineage, but not for other lineages [18], and the initiation of DBA pathology may occur because of disorder in energy usage switching during the differentiation of HSPCs into the erythroid lineage. Our results suggest that the disrupted OXPHOS pathway can lead to erythroid defects in patients with DBA, likely by interfering with energy switching during erythroid differentiation. These results demonstrate the potential for curing erythroid defects in DBA disease from the perspective of energy intervention.

Mitochondrial respiration can drive the commitment of the erythroid lineage at an early stage [30]. Our data showed that erythroid defects occurred at an early stage of erythroid differentiation, likely because HSCs that undergo metabolic reprogramming exit the quiescent state and enter the active state in response to the increased energy demand for differentiating from erythroid progenitors to Pro-erythroblasts (Pro-E). From the single-cell BM data, we observed that the OXPHOS pathway was preferentially active at the early stage of pro-E but not at the stage of erythroid progenitors, suggesting that this pathway defect can occur between HSPCs and pro-E, consistent with observations in patients with DBA [35]. This results in blockage at the early stage of erythroid differentiation, preventing cells from differentiating past the progenitor stage.

Defective heme biosynthesis is among the pathological mechanisms of DBA [10, 67]; both suppressed OXPHOS and heme biosynthesis pathways have been detected in patients with DBA. One possible explanation for defective heme biosynthesis in DBA cells is that the suppression of OXPHOS reduces the availability of ribosomes, causing an interference in the interaction between heme and ribosomes [68].

In DBA, certain mutations in ribosomal genes, including *RPL11, RPL5*, and *RPS19*, lead to ribosomal stress, which activates p53 and induces apoptosis [69]. Apoptotic cells were observed in established DBA cells; furthermore, erythroid progenitors or precursors are very sensitive to apoptosis caused by p53 activation [49, 70], and defective erythroid phenotypes can be rescued by reducing the expression of p53 [52, 71]. We identified a subset of apoptosis-related proteins and genes; however, p53 was not detected in the proteomic data. Apoptotic cells may be produced via defective ribosome biogenesis because of the specific roles of RPs in determining the erythroid lineage differentiation of HSPCs [35].

Previously, a single RP gene linked to ribosome biogenesis was extensively characterized [3, 12, 35]; however, the global defects in ribosome biogenesis that contribute to DBA pathology are largely unknown. Here, we revealed that suppression of OXPHOS in differentiating HSPCs leads to dysfunction of ribosomal biosynthesis, as demonstrated by the overall reduction in the expression of RPs, which may lead to abnormal translation of a selected erythroid-specific gene required for erythropoiesis [35, 44, 45]. Our results confirmed the relationship between an overall reduction in RPs and defects in erythropoiesis.

New drugs or treatments are needed for patients with DBA. Small-molecule compounds that stimulate erythropoiesis are a class of drugs with great potential for treating DBA. By enhancing the specific upregulation of ribosome biosynthesis and protein translation through the mTOR pathway, L-leucine supplementation dramatically improved anemia associated with RP loss in DBA in mouse and zebrafish models [8, 9, 72]. SMER28, a small-molecule inducer of the autophagy pathway that regulates erythropoiesis, was also identified as a potential therapeutic drug for DBA [54]. A DYRK inhibitor that interferes with the p53 pathway in erythroid progenitors was screened as a potential drug in a doxycycline-inducible mouse model for RPS19-deficient DBA [73]. Our results suggest that activation of the OXPHOS pathway can be targeted to improve erythroid defects in patients with DBA. Additionally, OXPHOS defects may occur because of insufficient electron transfer from complexes I and II to complex III, through which CoQ10 can largely alleviate erythroid defects at the base of disruption of the electronic transport chain, indicating its potential as a therapeutic drug for DBA. Thus, the OXPHOS pathway is a target of interest for DBA drug screening.

Our study has limitations. First, while we observed inhibition of OXPHOS in 7 of 10 DBA cases, the sample size was very small. Second, our study of the pathogenesis of DBA used an in vitro cell model; thus, the association of OXPHOS inhibition with erythroid defects should be confirmed in vivo and explored in higher resolution at the single cell level in the BM microenvironment of DBA. Finally, we were unable to determine the reason behind OXPHOS pathway disruption in some DBA cases. We propose that it can be caused by disorders in metabolic switching and alternative splicing events during the differentiation of HSCs into the erythroid lineage under abnormal conditions in BM.

In summary, we examined the relationship between the OXPHOS pathway and erythropoiesis and discovered that disruption of OXPHOS may lead to DBA. The disruption of OXPHOS is an alternative pathway of DBA pathogenesis and recapitulates the features in patients with DBA. We determined the molecular mechanism by which disruption of OXPHOS leads to DBA and reinforced the relationship between OXPHOS and ribosomal biogenesis in DBA. This study improves our understanding of erythropoiesis and provides insights into the pathology of DBA, which may improve translational research and treatments for this disease.

## Supporting information

Clinical information of patients

Primer sequences

Differentiate of erythroid progenitors (CD71+ GlyA-) in both BM and cord blood sources

Expression of ribosome-related genes

Defective phenotypes after the expression of RanGAP1

## Acknowledgements

This research was supported by the Strategic Priority Research Program of the Chinese Academy of Sciences (grant no. XDA16010602), National Natural Science Foundation of China (Grant Nos. 81870097, 82070114, and 81700116), and National Key Research and Development Program of China (Grant Nos. 2017YFC0907400, 2016YFC0901700, 2018YFC0910700, and 2017YFC0907403).

## Author contributions

XF, XZ, and ZZ conceived and supervised the study; RX, LZ, ZX, and QZ analyzed the data; ZZ and RX designed the experiments; RX, LZ, JZ, LZ, and JW performed the experiments; ZZ, RX, and SC drafted the manuscript; XC and YW helped with sample collection. XF, XZ, ZZ, and WY revised the manuscript. All authors have read and approved the final manuscript.

## Figure legends

**Supplemental Figure 1. Suppression of OXPHOS leads to defects in erythroid progenitor differentiation**

A–B. Flow cytometry analysis showing that the suppression of OXPHOS began decreasing in the percentage of erythroid progenitors/precursors (CD71+GlyA-) derived from Rot-treated HSPCs in healthy BM and human cord blood in differentiating DBA cells on day 3. C. Real-time PCR analysis revealing that the expression of some heme biosynthesis-related genes began decreasing in differentiating DBA cells on day 3. D. Detection of gene expression of erythroid-specific TFs and AHSP in differentiated DBA cells on day 3 using real-time PCR assay. E. Flow cytometry analysis showing increased apoptosis in differentiating DBA cells visualized on days 3 and 7. F. Real-time PCR analysis showing upregulated expression of apoptosis representative genes in differentiating DBA cells on days 3 and 7, particularly CDKN1A. ACTIN was used as a reference gene in real-time PCR assays. *p < 0.05; ***p < 0.001.

**Supplementary Figure 2. Suppression of OXPHOS leads to ribosomal defects in differentiating DBA cells**

A. Real-time PCR analysis showing significant overall reduction in the expression of most DBA causative genes identified in differentiating DBA cells on day 3. B. Real-time PCR analysis showing a decrease in gene expression of RPS3 in differentiating DBA cells on day 7. GAPDH was used as a reference gene in real-time PCR assays. *p < 0.05; ***p < 0.001. C. Immunofluorescence analysis of RPS3 showing defects in ribosomes in differentiating DBA cells on day 7. HSPCs were derived from human cord blood.

**Supplemental Figure 3. Knockdown of RanGAP1 leads to erythroid and ribosomal defects during terminal erythroid differentiation**

A. Real-time PCR analysis showing the knockdown of RanGAP1 by two independent shRNAs in differentiated HSPCs on day 14. B. Western blotting assay showing that the knockdown of RanGAP1 leads to a decrease in the expression of erythroid-specific proteins, including HBG and GATA1, on day 14. GAPDH was used as a loading control. C. Flow cytometry analysis showing a decrease in the number of CD71+GlyA+ double-positive erythroid cells in differentiated cells with RanGAP1 knockdown on day 14. D. Giemsa staining revealing a delay in the terminal erythroid differentiation of HSPCs with RanGAP1 knockdown on day 14. E. Real-time PCR analysis showing a decrease in the expression of globin genes in differentiated cells with RanGAP1 knockdown on day 14. F. Real-time PCR analysis showing a decrease in the expression of ribosomal representative DBA causative genes in differentiated cells with RanGAP1 knockdown on day 14. HSPCs were derived from human cord blood and GAPDH was used as a reference gene in real-time PCR assays. *p < 0.05; ***p < 0.001.

## References

1. van Dooijeweert, B., et al., Pediatric Diamond-Blackfan anemia in the Netherlands: An overview of clinical characteristics and underlying molecular defects. European Journal of Haematology, 2018. 100(2): p. 163–170.

2. Willig, T.N., H. Gazda, and C.A. Sieff, Diamond-Blackfan anemia. Current Opinion in Hematology, 2000. 7(2): p. 85–94.

3. Ulirsch, J.C., et al., The Genetic Landscape of Diamond-Blackfan Anemia. Am J Hum Genet, 2018. 103(6): p. 930–947.

4. Da Costa, L., T. Leblanc, and N. Mohandas, Diamond-Blackfan anemia. Blood, 2020. 136(11): p. 1262–1273.

5. Dutt, S., et al., Haploinsufficiency for ribosomal protein genes causes selective activation of p53 in human erythroid progenitor cells. Blood, 2011. 117(9): p. 2567–2576.

6. Yip, B.H., et al., Effects of L-leucine in 5q-syndrome and other RPS14-deficient erythroblasts. Leukemia, 2012. 26(9): p. 2154–2158.

7. Ludwig, L.S., et al., Altered translation of GATA1 in Diamond-Blackfan anemia. Nat Med, 2014. 20(7): p. 748–53.

8. Jaako, P., et al., Dietary L-leucine improves the anemia in a mouse model for Diamond-Blackfan anemia. Blood, 2012. 120(11): p. 2225–2228.

9. Payne, E.M., et al., L-leucine improves the anemia and developmental defects associated with Diamond-Blackfan anemia and del(5q) MDS by activating the mTOR pathway. Blood, 2012. 120(11): p. 2214–2224.

10. Yang, Z.T., et al., Delayed globin synthesis leads to excess heme and the macrocytic anemia of Diamond Blackfan anemia and del(5q) myelodysplastic syndrome. Science Translational Medicine, 2016. 8(338).

11. Mills, E.W. and R. Green, Ribosomopathies: There’s strength in numbers. Science, 2017. 358(6363).

12. Devlin, E.E., et al., A transgenic mouse model demonstrates a dominant negative effect of a point mutation in the RPS19 gene associated with Diamond-Blackfan anemia. Blood, 2010. 116(15): p. 2826–2835.

13. Torraco, A., et al., Mouse models of oxidative phosphorylation defects: Powerful tools to study the pathobiology of mitochondrial diseases. Biochimica Et Biophysica Acta-Molecular Cell Research, 2009. 1793(1): p. 171–180.

14. Steele, S.L., S.V. Prykhozhij, and J.N. Berman, Zebrafish as a model system for mitochondrial biology and diseases. Translational Research, 2014. 163(2): p. 79–98.

15. Chen, M.L., et al., Erythroid dysplasia, megaloblastic anemia, and impaired lymphopoiesis arising from mitochondrial dysfunction. Blood, 2009. 114(19): p. 4045–4053.

16. Cliff, T.S. and S. Dalton, Metabolic switching and cell fate decisions: implications for pluripotency, reprogramming and development. Current Opinion in Genetics & Development, 2017. 46: p. 44–49.

17. Oburoglu, L., et al., Metabolic regulation of hematopoietic stem cell commitment and erythroid differentiation. Current Opinion in Hematology, 2016. 23(3): p. 198–205.

18. Liu, X., et al., Regulation of mitochondrial biogenesis in erythropoiesis by mTORC1-mediated protein translation. Nature Cell Biology, 2017. 19(6): p. 626-+.

19. Guo, Y., et al., Transfer RNA detection by small RNA deep sequencing and disease association with myelodysplastic syndromes. Bmc Genomics, 2015. 16.

20. Vlachos, A., et al., Diagnosing and treating Diamond Blackfan anaemia: results of an international clinical consensus conference. British Journal of Haematology, 2008. 142(6): p. 859–876.

21. Boria, I., et al., The Ribosomal Basis of Diamond-Blackfan Anemia: Mutation and Database Update. Human Mutation, 2010. 31(12): p. 1269–1279.

22. Vlachos, A. and E. Muir, How I treat Diamond-Blackfan anemia. Blood, 2010. 116(19): p. 3715–3723.

23. Ren, Y., et al., Regulatory association of long noncoding RNAs and chromatin accessibility facilitates erythroid differentiation. Blood Adv, 2021.

24. Xin, Z.J., et al., Mapping Human Pluripotent Stem Cell-derived Erythroid Differentiation by Single-cell Transcriptome Analysis. Genomics Proteomics & Bioinformatics, 2021. 19(3): p. 358–376.

25. Hu, J.P., et al., Isolation and functional characterization of human erythroblasts at distinct stages: implications for understanding of normal and disordered erythropoiesis in vivo. Blood, 2013. 121(16): p. 3246–3253.

26. Li, J., et al., Isolation and transcriptome analyses of human erythroid progenitors: BFU-E and CFU-E. Blood, 2014. 124(24): p. 3636–3645.

27. Wen, Y.Y., et al., Quantitative proteomic analysis of scleras in guinea pig exposed to wavelength defocus. Journal of Proteomics, 2021. 243.

28. Narla, A. and B.L. Ebert, Ribosomopathies: human disorders of ribosome dysfunction. Blood, 2010. 115(16): p. 3196–3205.

29. Narla, A., S.N. Hurst, and B.L. Ebert, Ribosome defects in disorders of erythropoiesis. International Journal of Hematology, 2011. 93(2): p. 144–149.

30. Rossmann, M.P., et al., Cell-specific transcriptional control of mitochondrial metabolism by TIF1 gamma drives erythropoiesis. Science, 2021. 372(6543): p. 716-+.

31. Richard, A., et al., Erythroid differentiation displays a peak of energy consumption concomitant with glycolytic metabolism rearrangements. Plos One, 2019. 14(9).

32. Dianzani, I., E. Garelli, and U. Ramenghi, Diamond-Blackfan anemia: A congenital defect in erythropoiesis. Haematologica, 1996. 81(6): p. 560–572.

33. Sankaran, V.G., et al., Exome sequencing identifies GATA1 mutations resulting in Diamond-Blackfan anemia. Journal of Clinical Investigation, 2012. 122(7): p. 2439–2443.

34. Chennupati, V., et al., Ribonuclease inhibitor 1 regulates erythropoiesis by controlling GATA1 translation. J Clin Invest, 2018. 128(4): p. 1597–1614.

35. Khajuria, R.K., et al., Ribosome Levels Selectively Regulate Translation and Lineage Commitment in Human Hematopoiesis. Cell, 2018. 173(1): p. 90-+.

36. Choesmel, V., et al., Impaired ribosome biogenesis in Diamond-Blackfan anemia. Blood, 2007. 109(3): p. 1275–1283.

37. Rodriguez, M.S., C. Dargemont, and F. Stutz, Nuclear export of RNA. Biol Cell, 2004. 96(8): p. 639–55.

38. Dahlberg, J.E. and E. Lund, Functions of the GTPase Ran in RNA export from the nucleus. Curr Opin Cell Biol, 1998. 10(3): p. 400–8.

39. Ritterhoff, T., et al., The RanBP2/RanGAP1(star)SUMO1/Ubc9 SUMO E3 ligase is a disassembly machine for Crm1-dependent nuclear export complexes. Nature Communications, 2016. 7.

40. Bischoff, F.R., et al., Rangap1 Induced Gtpase Activity of Nuclear Ras-Related Ran. Proceedings of the National Academy of Sciences of the United States of America, 1994. 91(7): p. 2587–2591.

41. Joseph, J., et al., SUMO-1 targets RanGAP1 to kinetochores and mitotic spindles. J Cell Biol, 2002. 156(4): p. 595–602.

42. Mercurio, S., et al., The heme exporter Flvcr1 regulates expansion and differentiation of committed erythroid progenitors by controlling intracellular heme accumulation. Haematologica, 2015. 100(6): p. 720–729.

43. Testa, U., Apoptotic mechanisms in the control of erythropoiesis. Leukemia, 2004. 18(7): p. 1176–1199.

44. Li, D. and J.L. Wang, Ribosome heterogeneity in stem cells and development. Journal of Cell Biology, 2020. 219(6).

45. Flygare, J. and S. Karlsson, Diamond-blackfan anemia: erythropoiesis lost in translation. Blood, 2007. 109(8): p. 3152–3160.

46. Chen, J., et al., FAM122A Inhibits Erythroid Differentiation through GATA1. Stem Cell Reports, 2020. 15(3): p. 721–734.

47. Yang, G.H., et al., MicroRNAs Are Involved in Erythroid Differentiation Control. Journal of Cellular Biochemistry, 2009. 107(3): p. 548–556.

48. Sieff, C.A., et al., Pathogenesis of the erythroid failure in Diamond Blackfan anaemia. British Journal of Haematology, 2010. 148(4): p. 611–622.

49. Ohene-Abuakwa, Y., et al., Two-phase culture in Diamond Blackfan anemia: localization of erythroid defect. Blood, 2005. 105(2): p. 838–846.

50. Wan, Y., et al., Transcriptome analysis reveals a ribosome constituents disorder involved in the RPL5 downregulated zebrafish model of Diamond-Blackfan anemia. Bmc Medical Genomics, 2016. 9.

51. Zhang, Z.J., et al., Assessment of hematopoietic failure due to Rpl11 deficiency in a zebrafish model of Diamond-Blackfan anemia by deep sequencing. Bmc Genomics, 2013. 14.

52. Jia, Q., et al., Transcriptome Analysis of the Zebrafish Model of Diamond-Blackfan Anemia from RPS19 Deficiency via p53-Dependent and -Independent Pathways. Plos One, 2013. 8(8).

53. Song, B.F., et al., Systematic transcriptome analysis of the zebrafish model of diamond-blackfan anemia induced by RPS24 deficiency. Bmc Genomics, 2014. 15.

54. Doulatov, S., et al., Drug discovery for Diamond-Blackfan anemia using reprogrammed hematopoietic progenitors. Science Translational Medicine, 2017. 9(376).

55. Heijnen, H.F., et al., Ribosomal Protein Mutations Induce Autophagy through S6 Kinase Inhibition of the Insulin Pathway. Plos Genetics, 2014. 10(5).

56. Narla, A., et al., L-Leucine improves the anaemia in models of Diamond Blackfan anaemia and the 5q-syndrome in a TP53-independent way. British Journal of Haematology, 2014. 167(4): p. 524–528.

57. Shah, D.I., et al., Mitochondrial Atpif1 regulates haem synthesis in developing erythroblasts. Nature, 2012. 491(7425): p. 608–12.

58. Viphakone, N., et al., TREX exposes the RNA-binding domain of Nxf1 to enable mRNA export. Nature Communications, 2012. 3.

59. Slomovic, S., et al., Polyadenylation of ribosomal RNA in human cells. Nucleic Acids Research, 2006. 34(10): p. 2966–2975.

60. Harvey Lodish, A.B., Chris A. Kaiser, Molecular Cell Biology, in Molecular Cell Biology. 2016, W. H. Freeman: New York. p. 626.

61. Johnson, A.W., E. Lund, and J. Dahlberg, Nuclear export of ribosomal subunits. Trends in Biochemical Sciences, 2002. 27(11): p. 580–585.

62. Du, W., et al., SCO2 Mediates Oxidative Stress-Induced Glycolysis to Oxidative Phosphorylation Switch in Hematopoietic Stem Cells. Stem Cells, 2016. 34(4): p. 960–971.

63. Simsek, T., et al., The Distinct Metabolic Profile of Hematopoietic Stem Cells Reflects Their Location in a Hypoxic Niche. Cell Stem Cell, 2010. 7(3): p. 380–390.

64. Takubo, K., et al., Regulation of Glycolysis by Pdk Functions as a Metabolic Checkpoint for Cell Cycle Quiescence in Hematopoietic Stem Cells. Cell Stem Cell, 2013. 12(1): p. 49–61.

65. Baumann, K., Stem cells: A metabolic switch. Nat Rev Mol Cell Biol, 2013. 14(2): p. 64–5.

66. Cabon, L., et al., AIF loss deregulates hematopoiesis and reveals different adaptive metabolic responses in bone marrow cells and thymocytes. Cell Death and Differentiation, 2018. 25(5): p. 983–1001.

67. Mercurio, S., et al., Alteration of heme metabolism in a cellular model of Diamond-Blackfan anemia. European Journal of Haematology, 2016. 96(4): p. 367–374.

68. Chiabrando, D. and E. Tolosano, Diamond Blackfan Anemia at the Crossroad between Ribosome Biogenesis and Heme Metabolism. Adv Hematol, 2014. 2010(8): p.: 790632.

69. Lipton, J.M. and S.R. Ellis, Diamond-Blackfan anemia: diagnosis, treatment, and molecular pathogenesis. Hematol Oncol Clin North Am, 2009. 23(2): p. 261–82.

70. Tsai, P.H., S. Arkin, and J.M. Lipton, An intrinsic progenitor defect in Diamond-Blackfan anaemia. Br J Haematol, 1989. 73(1): p. 112–20.

71. Ellis, S.R., Nucleolar stress in Diamond Blackfan anemia pathophysiology. Biochim Biophys Acta, 2014. 1842(6): p. 765–8.

72. Stipanuk, M.H., Leucine and protein synthesis: mTOR and beyond. Nutrition Reviews, 2007. 65(3): p. 122–129.

73. Siva, K., et al., A Phenotypic Screening Assay Identifies Modulators of Diamond Blackfan Anemia. Slas Discovery, 2019. 24(3): p. 304–313.

