## Supplementary material for "Oxidative phosphorylation pathway disruption is an alternative pathological mechanism leading to Diamond-Blackfan anemia": Primer sequences

**Table2 qPCR primers**

| **Primers** | **Sequence（5' –> 3'）** |
|---|---|
| *HBB* F | AGGAGAAGTCTGCCGTTACTG |
| *HBB* R | CCGAGCACTTTCTTGCCATGA |
| *HBG* F | TTCACAGAGGAGGACAAGGCTAC |
| *HBG* R | GCAGAGGCAGAGGACAGGTT |
| *HBE* F | TTTGGAAACCTGTCGTCTCC |
| *HBE* R | GGCTTGAGGTTGTCCATGTT |
| *GLRX5* F | CTCCGACAAGGCATTAAAGACT |
| *GLRX5* R | GACCAATGAACTGCCGCTTC |
| *IBA57* F | GACCAATGAACTGCCGCTTC |
| *IBA57* R | CTGGAGCCCGTACAAGATGAC |
| *ISCA1* F | TAGGTGTAAAAGTTGGTGTCCGA |
| *ISCA1* R | ACAAGTCCCTTTGATGTTTGGG |
| *ISCU* F | GGGTCCCTTGACAAGACATCT |
| *ISCU* R | CCTTTCACCCATTCAGTGGCTA |
| *BACH1* F | CTCAGCCTTAATGACCAGCGG |
| *BACH1* R | GCCTACGATTCTTGAGTGGAAG |
| *PPOX* F | CTGGATTCGCTCCGTTCGAG |
| *PPOX* R | CCCACGTAGAGGAACCTGT |
| *SLC25a37* F | GATGGGGACAGCCGAGATG |
| *SLC25a37* R | ACCGGGTACATGACCGAGT |
| *ALAS1* F | CGCCGCTGCCCATTCTTAT |
| *ALAS1* R | TCTGTTGGACCTTGGCCTTAG |
| *ALAS2* F | CAGTTCCTGTTTGGTATTGGACG |
| *ALAS2* R | TGCCTTCTGCACAATCTTGCT |
| *GATA1* F | CTGTCCCCAATAGTGCTTATGG |
| *GATA1* R | GAATAGGCTGCTGAATTGAGGG |
| *KLF1* F | GGTTGCGGCAAGAGCTACA |
| *KLF1* R | GTCAGAGCGCGAAAAAGCAC |
| *NFE2* F | GCAGGAACAGGGTGATACAGC |
| *NFE2* R | GCAGCTCGGTGATGGACAT |
| *BCL11A* F | ACAAACGGAAACAATGCAATGG |
| *BCL11A* R | TTTCATCTCGATTGGTGAAGGG |
| *AHSP* F | GGATCTCATTTCCGCAGGATTG |
| *AHSP* R | CTGCTGCCTGTAATAGTTGATGT |
| *BCL2* F | GGTGGGGTCATGTGTGTGG |
| *BCL2* R | CGGTTCAGGTACTCAGTCATCC |
| *CDKN1A* F | TGTCCGTCAGAACCCATGC |
| *CDKN1A* R | AAAGTCGAAGTTCCATCGCTC |
| *RPL5* F | GCTCGGAAACGCTTGGTGATA |
| *RPL5* R | CCCTCTATACGGGCATAAGCAAT |
| *RPL9* F | GCACAGTTATCGTGAAGGGC |
| *RPL9* R | TTACCCCACCATTTGTCAACC |

| Primers | Sequence（5' –> 3'） |
|---|---|
| *RPL11* F | AAAGGTGCGGGAGTATGAGTT |
| *RPL11* R | TCCAGGCCGTAGATACCAATG |
| *RPL15* F | CCCACCCGGCCTGATAAAG |
| *RPL15* R | CACGGCGAACACGAATCCT |
| *RPL18* F | ATGTGCGGGTTCAGGAGGTA |
| *RPL18* R | CTGGTCGAAAGTGAGGATCTTG |
| *RPL26* F | GACTTCCGACCGAAGCAAGAA |
| *RPL26* R | TGCACCCGTTCAATGTAGATAAC |
| *RPL27* F | TGGCTGGAATTGACCGCTAC |
| *RPL27* R | CCTTGTGGGCATTAGGTGATTG |
| *RPL31* F | CTCGGGCACTCAAAGAGATTC |
| *RPL31* R | CGGATTCGGTATGGCACATTC |
| *RPL35* F | AGCTCTCTAAGATCCGAGTCG |
| *RPL35* R | GAACACGGGCAATGGATTTCC |
| *RPL35A* F | TTGAAGGTGTTTACGCCCGAG |
| *RPL35A* R | TGCTTCGGAATTTGGCACGA |
| *RPS7* F | GTGAAGCCCAATGGCGAGAA |
| *RPS7* R | TGAGGTCCGAGTTCATCTCCA |
| *RPS10* F | ATGTTGATGCCTAAGAAGAACCG |
| *RPS10* R | CGTAGCCTCGGGACTTGAGA |
| *RPS15* F | CCCGAGATGATCGGCCACTA |
| *RPS15* R | CCATGCTTTACGGGCTTGTAG |
| *RPS17* F | GTTCGCACCAAAACCGTGAAG |
| *RPS17* R | CGCTTGTTCGTGTGGAAGT |
| *RPS19* F | AAGCTGAAAGTCCCCGAATGG |
| *RPS19* R | AGTTCTCATCGTAGGGAGCAAG |
| *RPS24* F | ATGAACGACACCGTAACTATCCG |
| *RPS24* R | CCGAATTTCTGTCTTAGGCACTG |
| *RPS26* F | TAACTGTGCCCGATGCGTG |
| *RPS26* R | GCTCGCTTCAGAAATGTCCC |
| *RPS27* F | ATGCCTCTCGCAAAGGATCTC |
| *RPS27* R | TGAAGTAGGAATTGGGGCTCT |
| *RPS28* F | GACACGAGCCGATCCATCATC |
| *RPS28* R | TGACTCCAAAAGGGTGAGCAC |
| *RPS29* F | CGCTCTTGTCGTGTCTGTTCA |
| *RPS29* R | CCTTCGCGTACTGACGGAAA |
| *RPS6* F | TGGACGATGAACGCAAACTTC |
| *RPS6* R | TTCGGACCACATAACCCTTCC |
| *RPS3* F | CTGGAGTTGAGGTGCGAGTTA |
| *RPS3* R | ACAGCAGTCAGTTCCCGAATC |
| *RanGAP1* F | AACCGTCTGGAGAATGATGG |
| *RanGAP1* R | CGCAAGGTCTTCAAGGTCTC |

| **Primers** | **Sequence（5' –> 3'）** |
| --- | --- |
| *RanGAP1-sh1-F* | GATCCGAACTTGTCATTCTGTGAAATcttcctgtcagaATTTCACAGAATGACAAGTTCtttttg |
| *RanGAP1-sh1-R* | AATTCAAAAAGAACTTGTCATTCTGTGAAATTCTGACAGGAAGATTTCACAGAATGACAAGTTCG |
| *RanGAP1-sh2-F* | GATCCGCAGGAACTCAAGCTCAACAActtcctgtcagaTTGTTGAGCTTGAGTTCCTGCtttttg |
| *RanGAP1-sh2-R* | AATTCAAAAAGCAGGAACTCAAGCTCAACAATCTGACAGGAAGTTGTTGAGCTTGAGTTCCTGCG |
