## Supplementary figures and images for "Oxidative phosphorylation pathway disruption is an alternative pathological mechanism leading to Diamond-Blackfan anemia"

### Defective phenotypes after the expression of RanGAP1

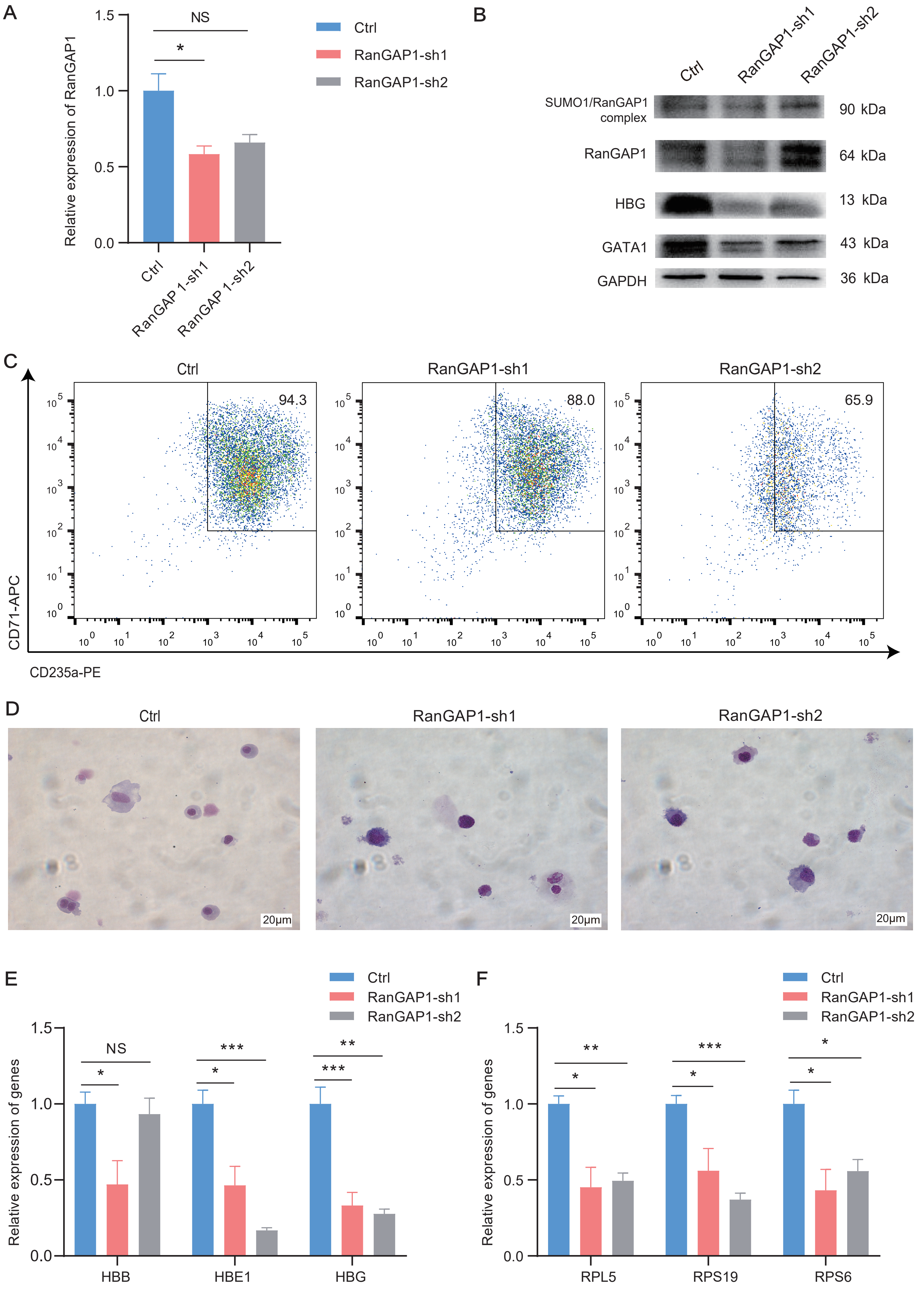

### Differentiate of erythroid progenitors (CD71+ GlyA-) in both BM and cord blood sources

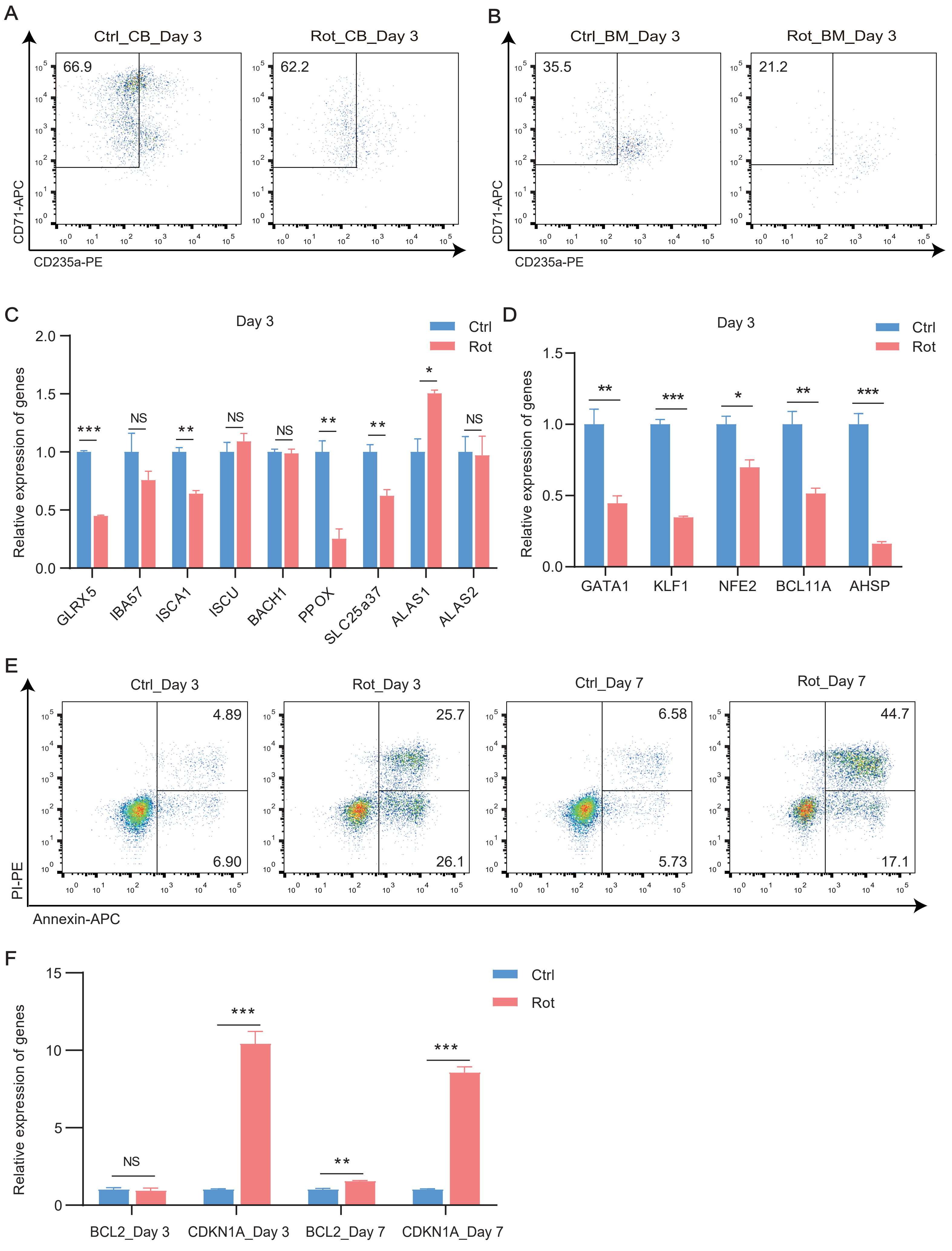

### Expression of ribosome-related genes

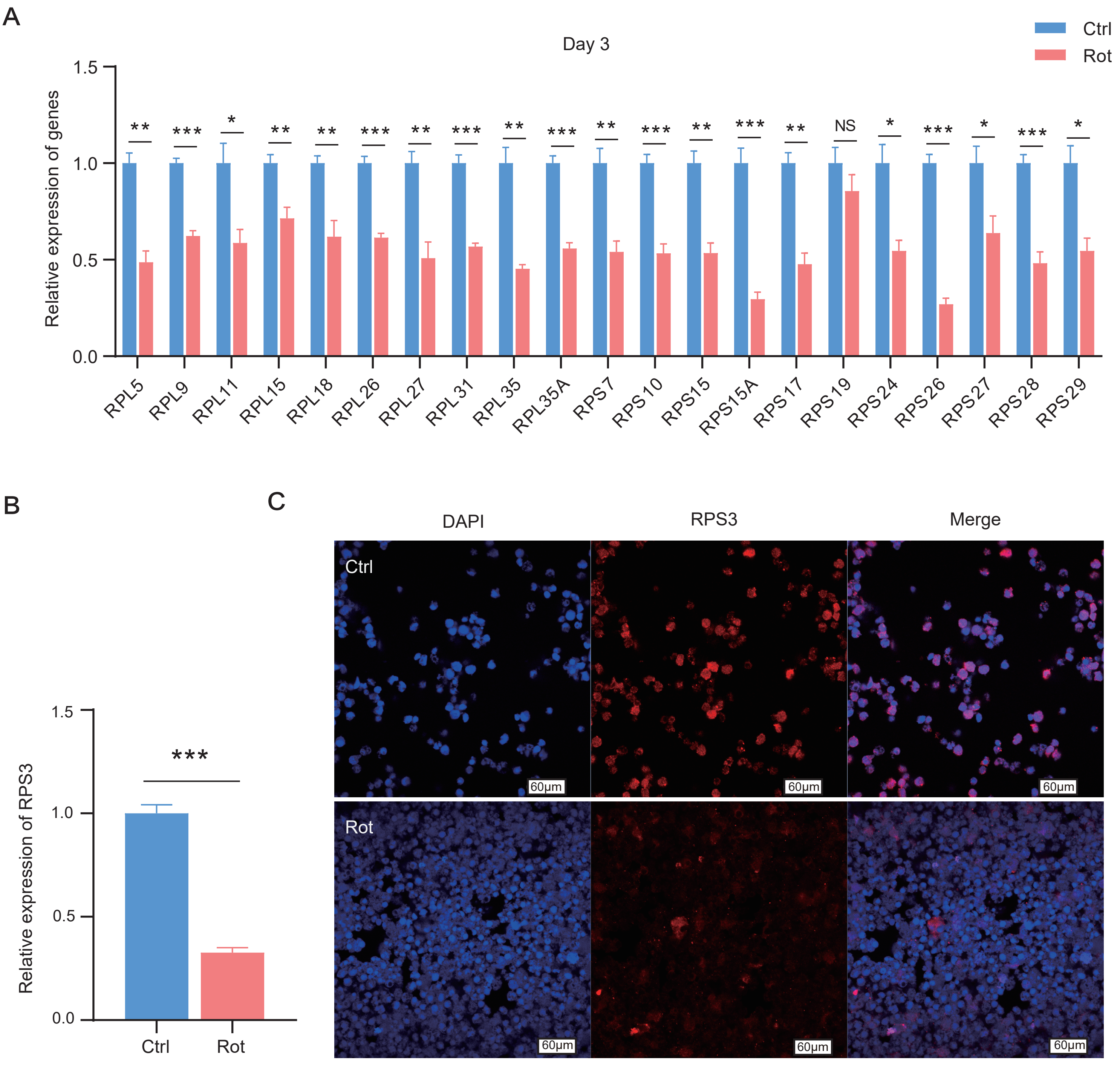
